# Comparison of commercially available differentiation media on morphology, function, and virus-host interaction in conditionally reprogrammed human bronchial epithelial cells

**DOI:** 10.1101/2023.04.12.536514

**Authors:** Nikhil T Awatade, Andrew T Reid, Kristy S Nichol, Kurtis F Budden, Punnam C Veerati, Prabuddha S Pathinayake, Christopher L Grainge, Philip M Hansbro, Peter AB Wark

**Affiliations:** School of Medicine and Public Health, University of Newcastle, Sydney, NSW, Australia; Immune Health, Hunter Medical Research Institute, Sydney, NSW, Australia; Asthma and Breathing, Hunter Medical Research Institute, Sydney, NSW, Australia; Centre for Inflammation, Centenary Institute and University of Technology Sydney, Sydney, NSW, Australia; Dept of Respiratory and Sleep Medicine, John Hunter Hospital

**Keywords:** epithelium, airways, air liquid interface, cell culture

## Abstract

**Introduction:** Primary air liquid interface (ALI) cultures of bronchial epithelial cells are used extensively to model airway responses. A recent advance is the development of conditional reprogramming that enhances proliferative capability. Several different media and protocols are utilized, yet even subtle differences may influence cellular responses. We compared the morphology and functional responses, including innate immune responses to rhinovirus infection in conditionally reprogrammed primary bronchial epithelial cells (pBECs) differentiated using two commonly used culture media.

**Methods:** pBECs from healthy participants (n = 5) were CR using γ-irradiated 3T3 fibroblasts and Rho Kinase inhibitor. CRpBECs were differentiated at ALI in either PneumaCult™ (PN-ALI) or Bronchial Epithelial Growth Medium (BEGM)-based differentiation media (BEBM:DMEM, 50:50, Lonza™) - (AB-ALI) for 28 days. Transepithelial electrical resistance (TEER), immunofluorescence, histology, cilia activity, ion channel function, and expression of cell markers were analyzed. Viral load was assessed by RT-qPCR and anti-viral factors quantified by Legendplex following Rhinovirus-A1b (RVA1b) infection.

**Results:** CRpBECs differentiated in PneumaCult™ were smaller and had a lower TEER and cilia beat frequency (CBF) compared to BEGM media. PneumaCult™ media cultures exhibited significantly increased *FOXJ1* expression, more ciliated cells with a larger active area, increased intracellular mucins, and increased calcium-activated chloride channel current. However, there were no significant changes in viral RNA or host antiviral responses.

**Conclusion:** There are distinct structural and functional differences in CRpBECs cultured in the two commonly used ALI differentiation media. Such factors need to be taken into consideration when designing and comparing CRpBECs ALI experiments.

## 1. Introduction

Air liquid interface (ALI) cultures of primary bronchial epithelial cells (pBECs) have proven to be an invaluable tool for studying respiratory diseases, providing a human cell model that recapitulates features of the airway epithelium *in vivo*^(1,2)^. The major limiting factors are the paucity of pBECs which can be obtained and cultured *ex vivo* from endobronchial brushing, as primary cells have a finite capacity for expansion and proliferation^(3,4)^. Conditional reprogramming (CR) is an increasingly popular technique to increase the number of pBECs available for investigation. Here, the concomitant use of irradiated feeder cells and rho kinase (ROCK) inhibitor have enabled large-scale expansion of pBECs, facilitating investigations of cell-cell interactions, morphology, epithelial cell function and interventions with limited starting cell numbers^(4–6)^.

While most laboratories have adopted a CR expansion method, several different media are available for the subsequent differentiation into ALI culture including Epi (Epithelix), EMM (PromoCell, Epithelix), mAir (PromoCell), bronchial epithelial growth medium (BEGM) based differentiation media (BEBM:DMEM, 50:50, Lonza) and PneumaCult ALI (STEMCELL Technologies). The constituents of most differentiation media are proprietary but are certainly distinct since the use of different differentiation media has been shown to affect stratification, cell phenotypes and the proportions of different cell types^(7)^. This can impact the consistency, reproducibility, and comparability of research findings. Several studies have examined differences between alternate epithelial cell differentiation media^(8–12)^, but, given the increasing prevalence of CR-expanded ALI cultures, it is now necessary to examine how these media may affect differentiation of CRpBECs, which, have not been investigated.

Of the few studies using CRpBECs, one used nasal epithelial cells and the other bronchial, though both assessed morphology, differentiation phenotype and assessed limited functional characteristics, leaving a gap in the understanding of the impact of differentiation media on these cells (studies are summarized in **Table 1**).

**Table 1:** Previous studies that compared different commercial differentiation media

| Publications | Expansion by CR | Differentiation Media | Bronchial/ Nasal/Tracheal Cultures | Thickness of cultures | Resistance | Cilia density | Cilia beat frequency | Ion channel function | Cell morphology (Cell size) | Mucosecretory markers (MUC5AC, goblet cells) | Virus infectivity | Inflammatory chemokines |
| --- | --- | --- | --- | --- | --- | --- | --- | --- | --- | --- | --- | --- |
| Current study | Yes (F media) | AB-ALI vs PN-ALI | Bronchial (adult) | Thicker in PN-ALI | Lower in PN-ALI | Higher density and FOXJ1 expression in PN-ALI | Higher in AB-ALI | Similar CFTR currents, higher CaCC currents | Smaller, more tightly packed cells in PN-ALI | Higher MUC5AC, but not significant in PN-ALI | More RSV infected cells in PN-ALI but not significant | Similar in both media |
| Livnat et al., 2023 (12) | No | MEM<br>USG-ALI<br>ALI from UNC<br>PN-ALI | CFBE41o Cells (bronchial) stably expressing cell line | - | - | - | - | Higher wt-CFTR currents in MEM media. | - | - | - | - |
| Luengen et al., 2020 (10) | No | mAir, PN-ALI, Epi, EMM | Nasal (donors between 10-40 years old) | - | Measured but not mentioned which media had better resistance | Increased ciliation in mAir media. Lower in PN-ALI and EMM | - | - | - | Higher MUC5AC in mAir cultures. PN-ALI, Epi and EMM cultures were negative for MUC5AC | - | - |
| Leung et al., 2020 (9) | No (BEBM - Lonza) | Comparison of AB-ALI vs PN-ALI | Bronchial (adult) | Thicker in PN-ALI | Lower in PN-ALI | Higher in PN-ALI | Similar in both media | - | - | Higher MUC5AC in PN-ALI | - | - |
| Broadbent et al., 2020 (8) | No | Promocell vs PN-ALI media | Nasal (paediatric) | - | Similar in both media | Increased ciliated cells in PN-ALI but not significant | - | - | Smaller, more tightly packed cells in PN-ALI | Similar MUC5AC in both media | More RSV infected cells in PN-ALI | - |
| Saint-Criq et al., 2020 (11) | Yes (F media) | Comparison of AB-ALI vs PN-ALI | Bronchial (adult) | Thicker in PN-ALI | Lower in PN-ALI | Higher in PN-ALI | - | Higher CFTR and baseline currents in PN-ALI | - | No difference in MUC5AC across both media | - | - |
| Ruiz García et al., 2019 (7) | No (BEBM - Lonza) | Comparison of AB-ALI vs PN-ALI | Nasal (adult) | - | - | Higher in PN-ALI | - | - | - | No detection of MUC5AC or SCGB1A1 in AB-ALI | - | - |
| Lee et al., 2019 (21) | Yes (F media) | Comparison of BEGM, PromoCell, LHC-8 and PN-ALI | Nasal (adult) | - | Lower in PN-ALI | Higher in BEGM | Lower in PN-ALI | - | - | Higher MUC5AC expression in PN-ALI | - | - |

**MEM:** minimum essential media, **USG:** Ultrosor-G (TM), **PN-ALI:** PneumaCult ALI, **50:50 BEBM and DMEM:** AB-ALI, **BEGM:** bronchial epithelial growth medium, **BEBM:** bronchial epithelial cell growth basal medium, **F media:**, **CaCC:** Calcium activated currents, **AECGM:** Promocell, **CRC:** conditionally reprogrammed culture # PN-Explus media to apical and basal side for initial 4 days on transwells, **mAir:** modified AECGM, **Epi:** MucilAir culture medium, **EMM:** EGM2-MucilAir-mixture, **wt-CFTR:** wild type CFTR, **MUC5AC:** Mucin 5AC.

Here, we compared the effects of two commonly used commercially available differentiation media in CRpBECs: PneumaCult™ and Bronchial Epithelial Growth Media (BEGM) based ALI media (Lonza™). Media were compared in terms of their impact on morphological features, epithelial barrier integrity, ciliary activity, ion channel function, mucus production, viral replication kinetics, and antiviral responses (96h post rhinovirus (RV)A1b infection) in CRpBECs. We found significant differences in morphological features and functional behavior (epithelial barrier function, ciliary activity, epithelial ion transport activity) between ALI cultures differentiated in PneumaCult and BEGM based media, but no significant difference in viral load or host antiviral responses after infection. These findings demonstrate how important the consideration of differentiation media is prior to the onset of culture-based experiments. This data highlights the impact that differing culture media have on overall CRpBECs morphology and emphasizes the importance of considering this when choosing the media for an individual study, and when comparing studies that have used different media.

## 2. Materials and Methods

### 2.1 Human pBECs procurement and processing

The study was approved by the Hunter New England Area Health Service Ethics Committee (05/08/10/3.09) and the University of Newcastle (Newcastle, NSW, Australia) Safety Committee (R5/2017). Human pBECs were obtained by endobronchial brushing during fibre-optic bronchoscopy (as previously described^(13)^ from healthy subjects *(n*=5) following written informed consent. All participants were non-smokers and had normal lung function with no evidence of respiratory disease in the preceding 4 weeks. Demographics of study participants are in **Table 2**.

**Table 2.**
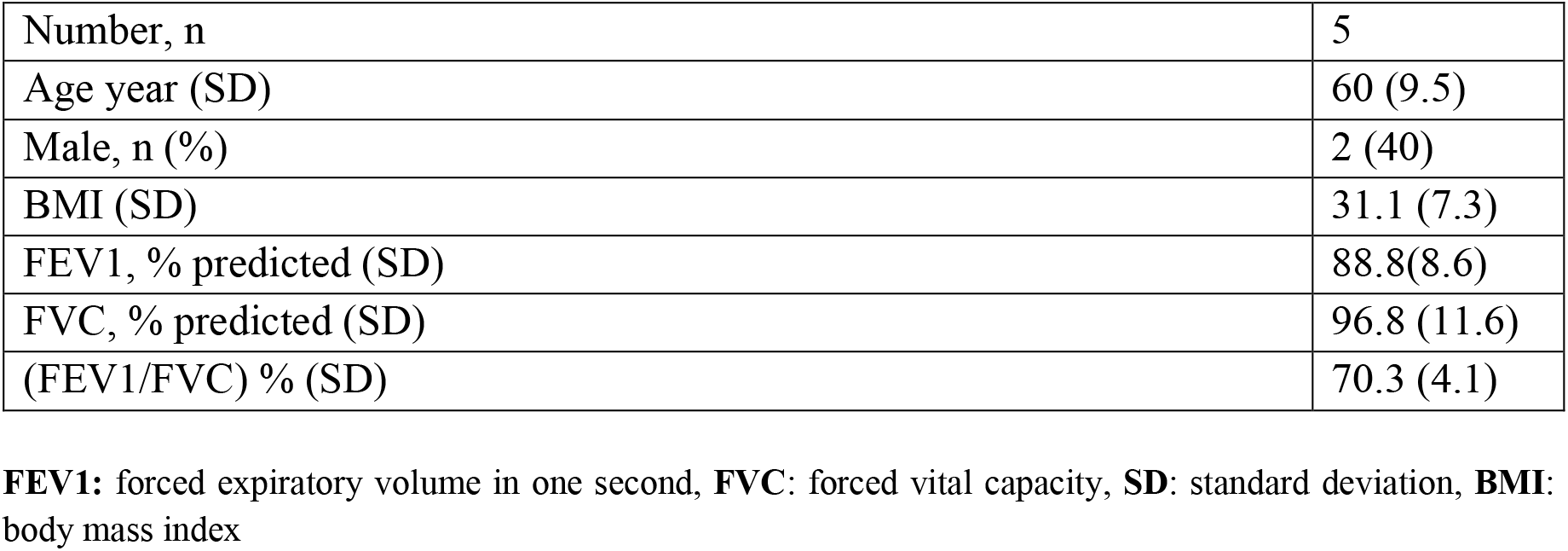
Clinical characteristics of healthy donors of primary broncho-epithelial cells

### 2.2 CR cell expansion culture/co-culture

HBE cells were previously established and expanded in standard BEGM Bulletkit (BEGM; Lonza). Passage 1 BEGM-expanded HBE cells were subsequently added to collagen I coated flasks and co-cultured with an equal amount of irradiated NIH/3T3 feeder cells in F-media containing the ROCK inhibitor Y-27632 as described previously^(5)^ (Table S1), with and without weaning steps. The weaning step was performed in three different stages, *i*) upon 50% confluency, media was switched to 70% Co-culture media: 30% BEGM complete media; *ii*) Two days later, media was changed to 50% Co-culture media:50% BEGM complete media; and *iii*) The next day cell media was changed to 50% Co-culture media without ROCK inhibitor:50% BEGM complete media. For cells that did not undergo the weaning step, media changes were performed every alternate day until 80-90% confluence with complete F-media containing ROCK inhibitor. Cells were passaged by differential trypsinization using a Trypsin/EDTA reagent pack (LONZA). Viability and cell count were assessed using the trypan-blue method.

### 2.3 NIH/3T3 feeder cell culture and irradiation

Cells of the NIH/3T3 mouse embryonic fibroblast line were cultured at 37°C, with 5% CO_2_ in DMEM (Sigma D5796) supplemented with 10% FBS and 1% (v/v) penicillin/streptomycin (Life Technologies, Australia). Cells at 80-90% confluency were trypsinized, and pelleted cells were resuspended in fresh culture media. For generation of a fibroblast ‘feeder’ layer, the NIH/3T3 cell suspension was exposed to 30 Gy γ-irradiation (RS 2000 X-Ray irradiator, RAD SOURCE) and then seeded into collagen I (PureCol; Advanced Biomatrix 5005) coated flasks at a 1:1 ratio with HBE cells, as previously described^(14,15)^.

### 2.4 ALI cultures

CR expanded HBE cells were seeded on collagen I coated 24-well Transwell membranes (Corning, USA; 6.5mm 0.4µm pore polyester membrane Sigma CLS3470) at a density of 1.5×10^5^ cells/insert. Cells were then cultured in two different differentiation media, bronchial epithelial base medium and Dulbecco’s modified eagle medium (BEBM:DMEM; 50:50) or PneumaCult Ex Plus/ALI (StemCell Technologies). In BEBM:DMEM 50:50 differentiation, for the initial 24 hours the cells were cultured in ALI initial media containing components listed in Table S2, for 3-5 days until confluent. Once confluent, apical media was removed (day 0 of ALI culture), basal media change was performed every second day with ALI final media that contains lower rhEGF concentrations (0.5 ng/mL), as previously described^(16)^. For PneumaCult differentiation, cells were cultured in PneumaCult Ex Plus for 4-5days until 100% confluent. Cells were then cultured at ALI by removing the growth medium from the apical surface and replacing the basal medium with PneumaCult ALI medium supplemented with hydrocortisone and heparin (StemCell technologies) according to the manufacturer’s instructions (Table S3), as previously described^(14)^. In both conditions, basal media was changed every second day (with corresponding media type) until day 28 of ALI culture. The apical surface was washed with phosphate-buffered saline (PBS, no Ca^2+^ and Mg^2+^) at 37°C once a week to remove excess mucus.

### 2.5 Transepithelial electrical resistance (TEER) measurements

TEER was measured at 7-day intervals from 7-28 days in ALI cultures using an Epithelial Tissue Voltohmmeter (EVOM2) and STX2 electrodes (World Precision Instruments, Sarasota, USA). The STX2 electrodes were equilibrated in D-PBS prior to measurement. Pre-warmed D-PBS (Lonza) was added to the apical side of the insert 5min prior to TEER measurement. Values were calculated after subtracting the blank value and according to the surface area of the inserts (0.33cm^2^). TEER is expressed as Ω.cm^2^. Results from each sample are the mean of 3-4 individual technical replicates.

### 2.6 Transepithelial ion transport assay

Ussing chamber measurements were performed for cultures with resistance values >200 Ω.cm^2^. Differentiated HBE ALI cultures (28-30 days old) were mounted in circulating Ussing chambers (Physiologic Instruments VCC MC8 multichannel voltage/current clamp). The epithelium was voltage-clamped, and the resistance and short-circuit current (I_sc_) were measured. The resistance of a filter and Ringer solutions in the absence of cells was subtracted from all measurements. Three-five transwells were analyzed per donor per condition. For I_sc_ recordings cells were bathed in 5ml of 37°C Krebs-bicarbonate-Ringer containing (mM): 115 NaCl, 25 NaHCO_3_, 2.4 K_2_HPO_4_, 0.4 KH_2_PO_4_, 1.2 CaCl_2_, 1.2 MgCl_2_ and 10 glucose, pH 7.4. Ringer solutions were continuously gassed with 95% O_2_- 5% CO_2_ and maintained at 37°C. After recording the stable baseline I_sc_ for 15min, cells were treated with pharmacological compounds (Table S4), in order: 100µM amiloride (apical) to inhibit epithelial sodium channel (ENaC)-mediated Na^+^ flux, 10µM forskolin and 100µM IBMX (apical and basal) to induce cAMP activation of the CFTR, 30µM CFTR_inh -172_ (apical) to inhibit CFTR-specific currents and 100µM ATP (apical) to activate calcium-activated chloride currents. Data were collected and analyzed using Acquire and Analyze software (v2.3, Physiologic Instruments). Results from each sample are presented as the mean of 10-15 individual technical replicates.

### 2.7 Cilia beating frequency and active area measurements

Cilia beating in differentiated CRpBEC cultures (frequency 3-18 Hz) was imaged on a Nikon eclipse Ti2 microscope (Nikon, Japan) and recorded with Video Savant 4.0 software using a high-speed digital video recorder. Recordings were made at 300 frames per second (fps) and a minimum of 512 frames were captured. Five fields of view were captured at random for each donor per differentiation media. Measurements of median cilia beat frequency were made using the CiliaFA plugin^(17)^. CiliaFA was used together with free open-source Fiji-ImageJ software (v1.53, ImageJ, US) and Microsoft Excel to perform analyses. Ciliary active area was calculated using Fiji-ImageJ software by performing built-in ‘Stack difference’ analysis to highlight areas of ciliary motion for each 512-frame file. Using the built-in Z-project plugin all 512 frames were ‘projected’ into 1 frame highlighting the regions of motile cilia. Thresholding was then applied identically to all projected images and active areas measured. Results from each sample are the mean of 5 fields of view.

### 2.9 Histology

ALI cultures at day 28 were washed with PBS, fixed in 10% neutral buffered formalin, embedded in paraffin and sectioned as described previously^(16)^. Alcian blue/periodic acid and Schiff (PAS) staining was performed also as previously described^(16)^. A minimum of 5 images were captured at random for each sample using a Nikon eclipse Ti2 microscope. Images were subjected to color deconvolution to quantify area of alcian blue/PAS overlap and were used in conjunction with measurements of total epithelial area to calculate the percentage of total epithelial area stained as previously described^(16)^.

### 2.10 Whole-mount immunolabelling and fluorescent microscopy

For immunolabelling of mucociliary differentiation markers, 28-day ALI cultures were first fixed in 4% paraformaldehyde for 15min followed by storage in PBS containing 50mM Glycine at 4°C as previously described^(16)^. Whole mount membranes were permeabilized with 0.1% v/v Triton-X 100 (Sigma) and blocked with 10% v/v Goat serum (Genesearch, Aus) in PBS. Membranes were divided prior to antibody exposure. Antibodies against acetylated tubulin (T7451, Sigma) and ZO-1 (33-9100, Invitrogen) were incubated overnight at 4°C. Membranes were washed in PBS and incubated with fluorescent secondary antibody Goat anti-mouse Alexafluor 594 (8890, Cell signaling technology, USA). Membranes were mounted on slides with Fluoromount-G containing DAPI (00-4959-52, Invitrogen, USA) placed under coverslips and sealed. A minimum of 3 images were captured per donor at random using Nikon Eclipse DS-Qi2 fitted with CoolLED box (pE-300). Images were then processed using ImageJ software (National Institute of Health, Bethesda, MD).

### 2.11 Viral infection

CRpBECs cultures were inoculated apically with human RVA1b at an MOI of 0.1 for 2h at 37°C or media control. The apical inoculum was removed, and cultures were washed 2 times with PBS. Apical and basal media samples were collected at 24, 48, and 96h post-inoculation and stored at -80 °C for cytometric bead array. Cells from the inserts were lysed and collected into RLT buffer containing B-mercaptoethnaol (QIAGEN) for downstream molecular analyses by RT-qPCR.

### 2.12 RNA extraction, cDNA synthesis, and quantitative PCR

Total RNA was extracted from ALI cultured cells lysed with RLT buffer containing β- mercaptoethnaol (QIAGEN) and frozen at -80°C for downstream analysis. RNA was extracted using RNeasy Mini Kits (QIAGEN, Germany) according to the manufacturer’s instructions. RNA quality and quantity were measured using a nano-drop 2000 spectrophotometer (Thermo Scientific™). RNA (200ng) was reverse transcribed to cDNA using high-capacity cDNA reverse transcription kits (Thermo Scientific™). qPCR was performed using a Quanstudio™ 6 as per the manufacturer’s instructions using TaqMan^®^ gene expression assays (ThermoFisher Scientific, Australia) and normalized to the ribosomal RNA (18s) housekeeping gene (Table S5). Relative gene expression was calculated using the 2^**-**ΔΔCt^ (where Ct is the threshold cycle) method, as previously described^(18–20)^.

### 2.13 Multiplex protein assessment

Apical supernatant from cells at each harvest time point was assessed using LEGENDPlex Human Anti-Virus Response Panel (BioLegend, San Diego, CA, USA), as per the manufacturer’s instructions and using a FACSCanto II flow cytometer (BD Biosciences, USA), as described previously^(20)^. Anti-viral interferons (IFN-β, λ1 and λ2/3) and inflammatory cytokines interleukins (IL-1β, 6), tumor necrosis factor-α (TNF-α), and interferon (IFN)-γ-inducible protein (CXCL10/IP-10) were measured.

### 2.14 Statistical Analysis

Data for resistance measures, cilia activity and Ussing chamber analyses are represented as dot plots with mean ± standard error of the mean (SEM). Data for all other measures are represented as with mean ± standard deviation (SD), with the respective number of experiments given in each figure legend. For statistical analysis, paired Student’s *t*-test or ANOVA were used as appropriate and data assessed with GraphPad Prism software (v9.3.1, San Diego, CA). A *p*-value <0.05 was considered statistically significant.

## 3. RESULTS

### 3.1 AB-ALI cultures have greater barrier function than PN-ALI cultures

The effect of differentiation media on epithelial barrier integrity was assessed by measuring the TEER of the cultures. Resistance values varied among donors in both culture media, with a range of 190-800 Ω.cm^2^. TEER data for both differentiation media displayed a general upward trend from day 7-28 post-ALI; with AB-ALI cultures increasing from 348±12.45 to 388.1±11.60 Ω.cm^2^ and PN-ALI cultures increasing from 221.8±6.92 to 301±19.49 Ω.cm^2^. At d7 and d14 time points AB-ALI had significantly higher TEER than PN-ALI. At d28 the TEER of AB-ALI cultures were higher though not significantly more than those of PN-ALI (388.1**±**11.60 *vs*. 301±19.49 Ω.cm^2^, respectively) (**Figure 1A**). These results suggest that differentiation media impacts epithelial barrier integrity, with AB-ALI cultures showing greater barrier function than PN-ALI cultures.

**Figure 1.**
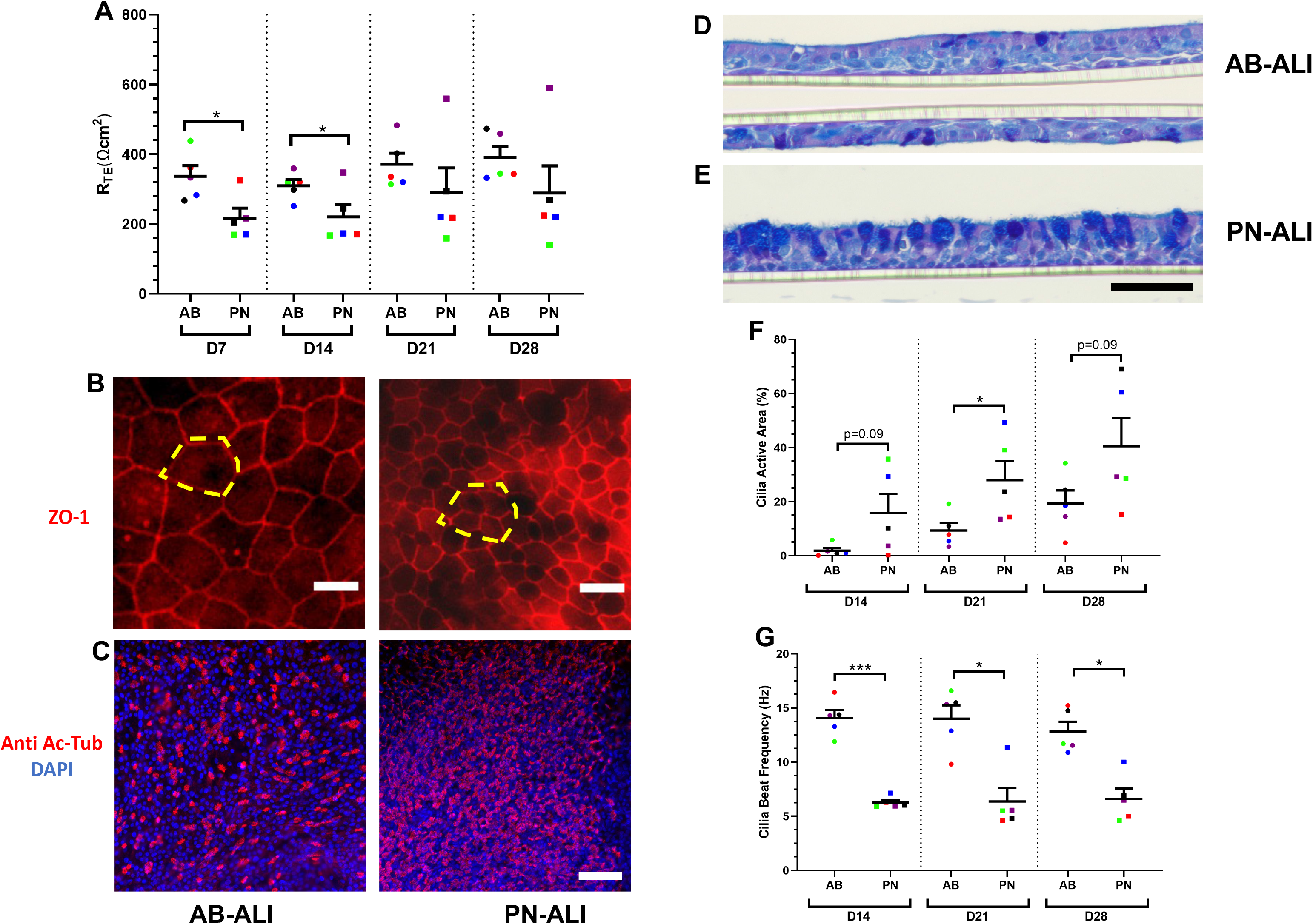
Structural and morphological characterization of AB-ALI and PN-ALI differentiated CRpBECs. (A) Trans-epithelial electrical resistance (TEER, R_TE_) values of AB-ALI and PN-ALI cultures. Each TEER data point represents an average of 10-15 transwells from each donor during the differentiation phase (d7-28). (B) Representative images of immunofluorescence staining of tight junctions (ZO-1, red). Yellow highlighted region shows 1 AB-ALI cell is equivalent to approximately 4 PN-ALI cells (Scale bar = 15μm). (C) Representative images of immunofluorescence staining of ciliated cells (anti-acetylated α −tubulin red), and DAPI stained nucleated cells (blue) (Scale bar = 40μm). (D, E) Representative images of Alcian blue/Periodic acid-Schiff-stained ALI cultures (Scale bar = 50μm). (F) Active cilia area and (G) cilia beat frequency measurements in AB-ALI and PN-ALI cultures. Five different fields of view were sampled per donor. A color-matched symbol in both groups represents an individual donor. n=5. Data presented as mean ± SEM. Data were analyzed using a paired Student’s *t*-test, *p* <0.05 was considered significant.

### 3.2 PN-ALI differentiated CRpBECs are smaller and more ciliated than with AB-ALI media

Following CRpBECs differentiation, we observed several morphological differences between PN-ALI and AB-ALI cultures. Immunofluorescent localization of ZO-1 (**Figure 1B**) revealed a considerable reduction in cell circumference/area when cultures were maintained in PN-ALI compared to AB-ALI. Indeed, these PN-ALI cells achieved only ∼25% the area of their AB-ALI counterparts. Similar staining thickness and uniformity was observed for ZO-1 labelling between the two media. A much higher proportion of cells exhibited anti-acetylated tubulin fluorescent labelling in PN-ALI cultures compared to AB-ALI, reflecting an increase in the proportion of ciliated cells with the former (**Figure 1C**). Longitudinal sections from each culture medium were cut and used for alcian blue periodic acid Schiff staining of mucus. Cells from cultures grown under PN-ALI conditions were variable in their overall mucin content, some matching closely to cultures grown under AB-ALI conditions while others had dense labelling for overlapping alcian blue, and PAS present in almost every ciliated-columnar cell. In contrast, AB-ALI cultures exhibited more consistent alcian blue/PAS staining that was present only in ∼ 5-10% of columnar cells for all donor samples. **Figure 1D, E**).

### 3.3 PN-ALI differentiated CRpBECs have elevated numbers of motile cilia but decreased cilia beat than AB-ALI cultures

The effect of differentiation media on ciliary function was assessed for two parameters: active ciliated area and, cilia beating frequency (CBF). Cilia motility was assessed on d14, 21, and 28 post-ALI, and beating cilia were observed from d14. PN-ALI cultures had significantly more ciliated cells with even coverage compared to AB-ALI, which showed sparsely distributed ciliated cells. The total active area of PN-ALI differentiated cultures were approximately double that of AB-ALI cultures at all time points, and became statistically significant at d21 (d14, d28; p=0.09). The greatest difference in total active area was observed at d28 (40.52±10.30 *vs* 19.27±4.9 %; *p* = 0.09) (**Figure 1F**). The CBF data for both differentiation media exhibited stable values from day d14-28. When compared to PN-ALI, CBF values for AB-ALI were ∼ two-fold higher and statistically significantly greater (d14 14.06±0.74 *vs* 6.26±0.23 Hz; *p* < 0.001, **Figure 1G**).

### 3.4 PN-ALI differentiated CRpBECs have higher baseline and calcium-activated chloride currents than AB-ALI cultures

The effect of differentiation media on ion transport was investigated by measuring transepithelial transport of amiloride-sensitive epithelial sodium channel (ENaC), forskolin-stimulated and CFTR_inh-172_-inhibited CFTR chloride secretion, and ATP-activated calcium-activated chloride channel (CaCC) currents (**Figure 2A**). All ion channel measurements showed donor-to-donor variability (**Table S6-7**). Baseline currents were higher but not significantly so in PN-ALI than AB-ALI differentiated cultures (33.57±9.54 *vs* 20.40±1.17 µA/cm^2^; *p*=0.0142, **Figure 2B**). The inhibition of ENaC currents by amiloride appeared to be greater in PN-ALI than in AB-ALI cultures, though the difference was not statistically significant (ΔI_sc-amiloride_ -11.10±1.98 *vs* -8.11±1.61 µA/cm^2^, **Figure 2C**). CFTR mediated chloride secretion was assessed by cAMP agonist forskolin and a CFTR-specific inhibitor (CFTR_inh-172_) was used to ensure that the currents were mediated by the CFTR chloride channel, and it completely inhibited the forskolin-induced currents in both cultures. No difference in forskolin-induced CFTR currents was observed between AB-ALI and PN-ALI cultures (ΔI_sc-fsk_ 12.23±1.79 *vs* 12.43±3.49 µA/cm^2^, **Figure 2D**). Although the difference was not statistically significant, PN-ALI cultures tended to have higher CFTR_inh-172_ currents than AB-ALI cultures (ΔI_sc-CFTRinh-172_ -28.87±8.55 *vs* -19.00±2.76 µA/cm^2^, **Figure 2E**). The addition of ATP at the end of the experiment resulted in transient CaCC currents. CaCC currents for PN-ALI were three times higher and statistically significantly higher than AB-ALI cultures (ΔI_sc-ATP_, 6.59±1.06 *vs* 2.29±0.59, µA/cm^2^; *p*<0.0001, **Figure 2F**). Overall, differentiation media affects ion transport with trends to differences in CFTR chloride and ENaC currents and large differences in CaCC currents.

**Figure 2.**
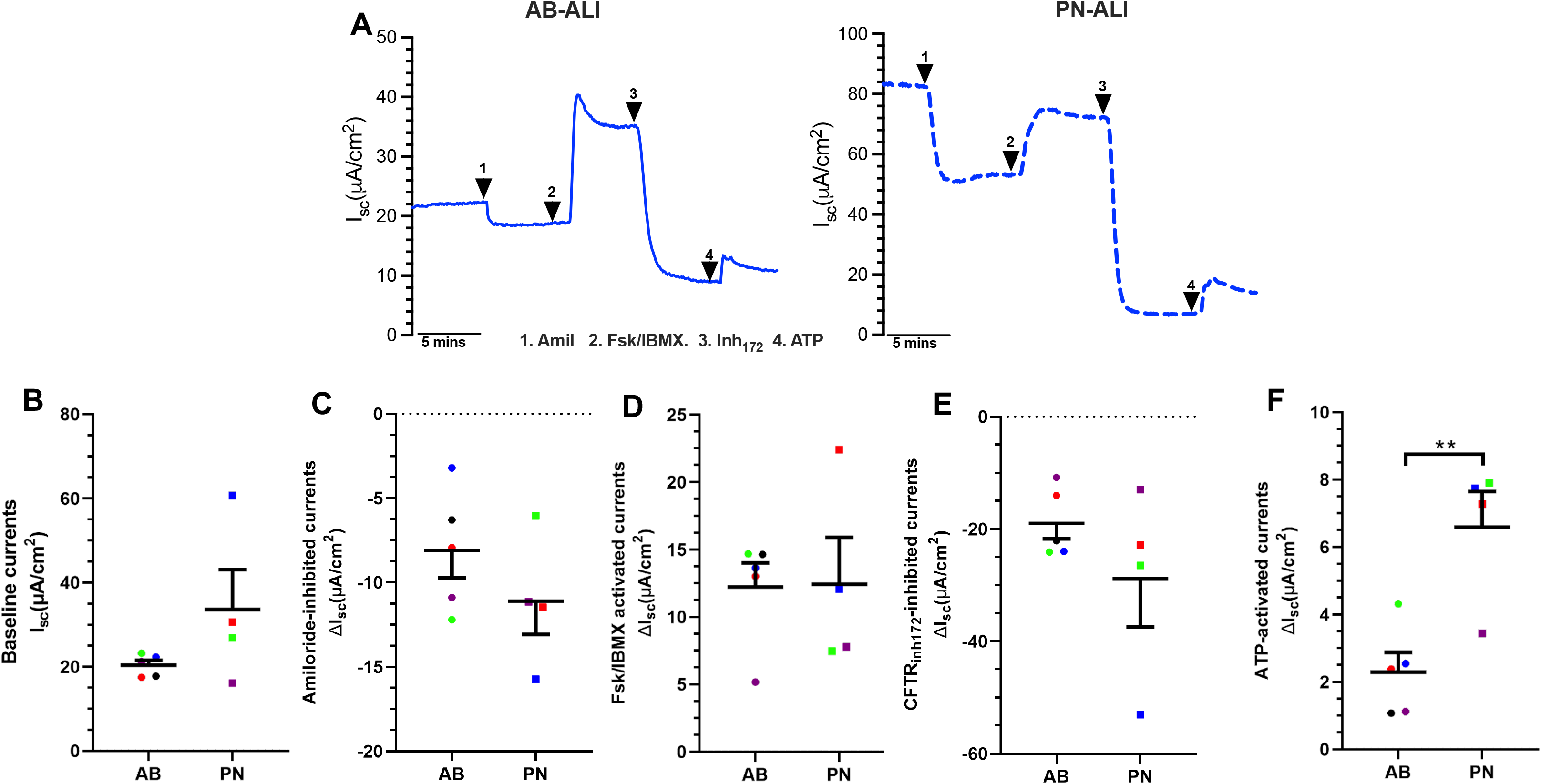
Ion transport measurements in AB-ALI and PN-ALI differentiated CRpBECs. (**A**) Representative Ussing chamber short circuit current (I_sc_) tracings recorded at 37°C for AB-ALI (blue line) and PN-ALI (dashed blue line) cultures. Mean values of (**B**) baseline short circuit currents (ΔI_sc_), (**C**) Amiloride inhibited ENaC currents, (**D**) Forskolin/IBMX activated CFTR, (**E**) CFTR_inh-172_ inhibited CFTR and (**F**) ATP-activated CaCC currents. n=5 for AB-ALI and n=4 for PN-ALI. Data presented as mean ± SEM. A color-matched symbol in both groups represents an individual donor. Data were analyzed using a paired Student’s *t*-test, *p* <0.05 was considered significant.

### 3.5 PN-ALI differentiated CRpBECs have increased FOXJ1 expression than AB-ALI cultures

Cell markers were assessed by qPCR. There were no differences in the expression of the secretory cell markers SPDEF (**Figure 3A**) enriched in goblet cells, or SGB1A1 (**Figure 3B**) enriched in club cells. However, expression of FOXJ1 (**Figure 3C**), a marker of ciliogenesis, was significantly increased in PN-ALI compared to AB-ALI cultures.

**Figure 3.**
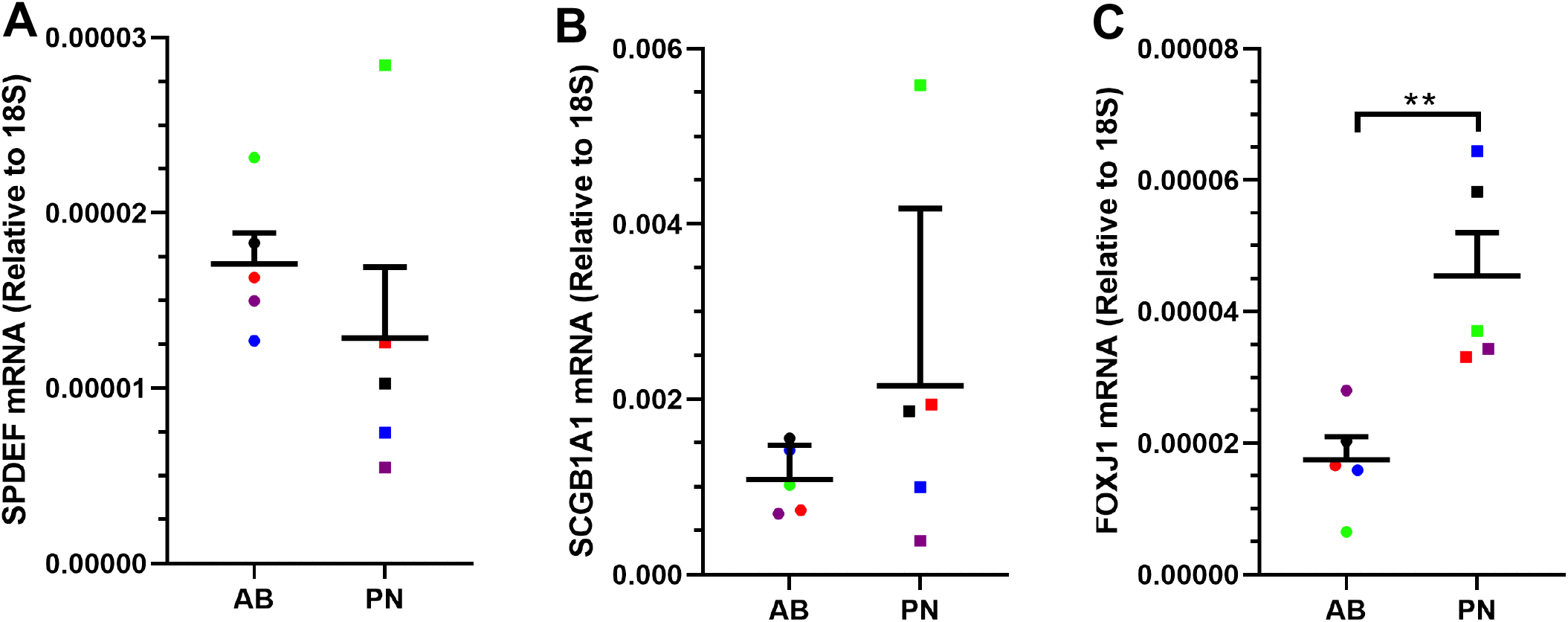
Cell marker gene expression in AB-ALI and PN-ALI differentiated CRpBECs. mRNA levels were measured by real-time quantitative PCR (RT-qPCR). Gene expression for (**A**) SPDEF, (**B**) SCGB1A1, (**C**) FOXJ1 is reported as [fold change (2^ddct)]. Data presented as mean ± SD. A color-matched symbol in both groups represents an individual donor. Data were analyzed using a paired Student’s *t*-test, *p*<0.05 was considered significant.

### 3.6 Response to RV infection, viral load and cytokine production did not differ between PN-ALI and AB-ALI

Following infection with RVA1b, viral load was assessed by qPCR and anti-viral mediators in the apical supernatant assessed at 24, 48 and 96 hours (**Figure S1**). There was considerable variability between individual cells. Viral load ranged from 1.56×10^5^ to 4.16×10^7^ copies/µL cDNA and was similar at all three timepoints. There were no differences between PN-ALI and AB-ALI cultures at any timepoint (**Figure 4A**). Production of the interferons IFN-β and IFN-λ was induced by RVA1b infection after 48 and 96 hours, though only reached significance in AB-ALI cells (**Figure 4 B, C**). Similarly, IP-10 appeared to be induced at all time points, though this was only statistically significant in AB-ALI at 48 and 96 hours (**Figure 4D**). The abundance of IP-10 after 48 hours was significantly greater than at 96 hours. Whilst there was a trend towards lower IP-10 production in PN-ALI after infection, there was no difference in interferon/cytokine abundance between PN-ALI and AB-ALI at any timepoint.

**Figure 4.**
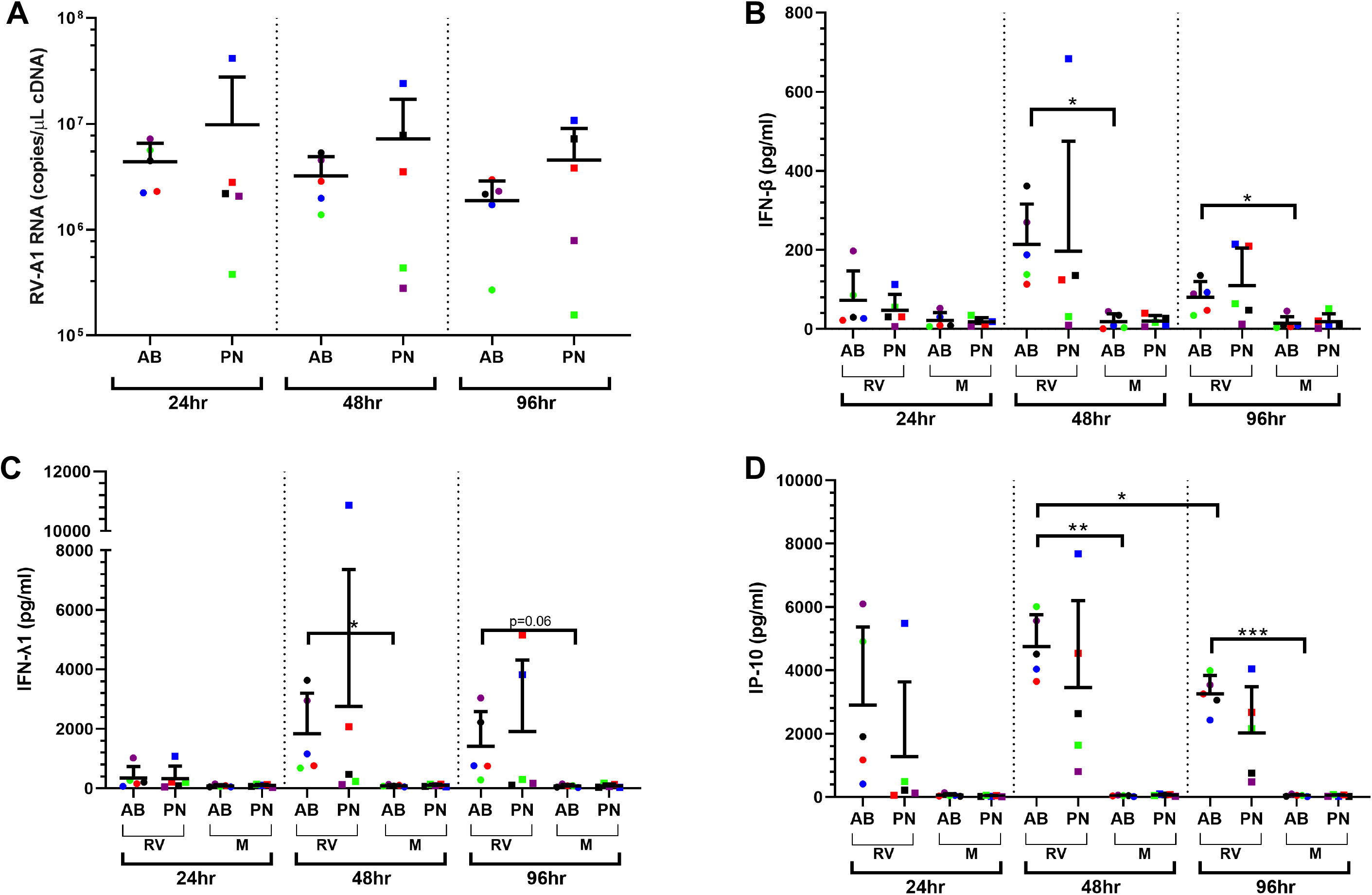
Viral load and cytokine production between AB-ALI and PN-ALI CRpBECs grown at air liquid interface and infected with rhinovirus RV-A1. (**A**) Viral RNA was quantified over time using real-time quantitative PCR (RT-qPCR). A multiplex protein quantification assay (LEGENDPlex) was performed using apical supernatant to determine changes in (**B**) IFN-β, (**C**) IFN-λ1, and (**D**) IP-10. Data presented as mean ± SD. A color-matched symbol in both groups represents an individual donor. Data were analyzed using a paired one-way ANOVA with Holm-Šídák pos hoc test. *p*<0.05 was considered significant.

## 4. DISCUSSION

The airway epithelium is a critical barrier that plays a crucial role during infection and is often dysregulated in respiratory diseases such as asthma, chronic obstructive pulmonary disease (COPD), and cystic fibrosis (CF). Differentiated cultures of human pBECs are a valuable tool for investigating airway epithelial responses. However, only a limited number of pBECs can be obtained from human donors, with their capacity for expansion and proliferation restricted, making it challenging to study their function and response to interventions. The recent development of CR technology allows for large-scale expansion of pBECs, which can be used for larger and more complex studies. Combining the CR expansion method with ALI culture creates a powerful tool for the study of respiratory diseases, however, varying culture conditions including the choice of media may introduce variability in results. Several studies have compared a small number of parameters for different CR ALI culture media (**Table 1**) but have not compared differences in the effects of differentiation media. Here, we provide a detailed characterization of two widely used and well reported differentiation media for CR ALI cultures, characterizing a wide range of parameters to identify morphological, functional and immune phenotypes. We show that PneumaCult™ and BEGM-based media result in significant differences in barrier function, active ciliated area/mucociliary function, cilia beating frequency, and ion transport. These differing effects need to be considered when reporting results, especially for assessments of morphology and studies of mucociliary clearance. In contrast, we found no differences in TEER and only minor differences in responses to infection with RVA1b in terms of viral load and antiviral responses.

The formation of tight junctions is crucial for the establishment of a functional epithelial barrier, and high TEER values are indicative of well-differentiated and tightly connected epithelium. Following differentiation, we found that both media yielded robust TEER values, suggesting that neither compromised epithelial tight junction formation. This is important for electrophysiological studies that rely on the integrity of the epithelial barrier. Interestingly, we observed that PN-ALI differentiated epithelia had a lower TEER compared to AB-ALI differentiated epithelia, consistent with previously published studies and likely attributable to a combination of reduced expression of sealing claudins and increased electrogenic ion transport^(9,11,21)^. Conversely, the higher TEER in AB-ALI differentiated cultures could be due to a squamous epithelium phenotype, which has been shown to confer higher TEER values^(9,22)^.

Significant differences in markers of mucociliary differentiation were observed between PN-ALI and AB-ALI most noticeably with the distribution of ciliated cells at the apical surface. PN-ALI cultures exhibited a relatively even covering of ciliated cells compared to the sparse distribution in AB-ALI. We also observed significantly increased expression of FOXJ1 mRNA, a marker of ciliogenesis, in PN-ALI compared to AB-ALI cultures, which was consistent with prior findings^(7–9)^. We previously reported that CR of cells from asthmatic donors leads to lower expression of FOXJ1 and may suggest that differentiation in PN-ALI mitigates this effect^(23)^. Interestingly, although there was a greater area covered by cilia the CBF was significantly lower in PN-ALI compared to AB-ALI cultures. Given that the PN-ALI media was associated with a non-significant decrease in the goblet cell marker SPDEF and an increase in the club cell marker SCGB1A1, we speculate that differences in secretory function may contribute to more viscous mucus produced by PN-ALI differentiated cells, that may prevent cilia from beating as freely. However, prior studies by Ruiz Garcia et al., 2019, identified SCGB1A1 as being associated with goblet cells in PneumaCult ALI media^(7)^. Further studies are needed to fully characterize secretory cell distribution and function and understand the underlying mechanisms of these differences.

For measurements of transepithelial ion transport using the Ussing chamber, PN-ALI cultures exhibited higher baseline currents most likely due to the small size and greater number of cells increasing the total surface area of the basolateral and apical membrane, as reported previously^(22)^. Moreover, PN-ALI cultures had higher though not significant differences in amiloride inhibited ENaC, this may be due to higher baseline levels of the function of these channels leading to an increase in overall ENaC conduction, as previously reported^(24)^. We did not observe any differences in CFTR conduction. We did also observe increased CaCC in PN-ALI cultures compared to AB-ALI. Although the mechanism is unclear, this may be due to increased electrogenic ion transport as suggested previously^(11)^ or higher expression of CaCC or more intracellular levels of Ca^2+^ in PN-ALI cultures.

Both media resulted in cultures that were successfully infected with RV with almost similar viral growth kinetics in line with the previous findings^(8)^. This and only other study compared the effect of differentiation media on viral growth kinetics or exogenous challenge such as infection. In agreement with the previous findings, we also observed higher though not significantly different levels of IFN- λ1 secreted from RV-infected PN-ALI cultures. Broadbent et al proposed that this may be due to the higher cell density in the cultures^(8)^. The process of conditional reprogramming has been shown to impact viral load, growth kinetics, and immune responses but our findings suggest that the selection of differentiation media has little additional impact^(23)^. However, this should be validated for other differentiation media before mainstream use.

Our findings demonstrate that the choice of culture differentiation media can impact mucociliary differentiation, with alterations in epithelial ion channel function in CRpBECs, which is likely to result in important differences in these features when comparing results from different experiments. These observations have important implications for researchers working with *in vitro* models of the airway epithelium for disease modelling and highlight the need for a better understanding of the factors that influence mucociliary differentiation in ALI cultures. Furthermore, efforts should be made to establish standardized differentiation protocols for ALI cultures as well as to improve the accessibility of information regarding commercially available differentiation media.

## Author contributions

NTA, KSN and ATR conceived and designed research, NTA, KSN and ATR performed experiments, NTA, KSN, KFB and ATR analyzed data; NTA, KSN, ATR, KFB, CLG, PMH and PABW interpreted results of experiments, NTA, KFB, KSN, ATR, PCV, and PSP prepared figures; NTA, KFB, and ATR drafted manuscript; NTA, KFB, PMH and PABW edited and revised manuscript; NTA, KSN, ATR, KFB, PCV, PSP, CLG, PMH and PABW approved final version of manuscript.

## Data availability statement

The raw data supporting the conclusion of this article will be made available by the authors, without undue reservation.

## Supplementary tables

**Supplementary Table S1.** Components of CR cell expansion media.

| Components | Concentration | Supplier | Catalogue number |
| --- | --- | --- | --- |
| DMEM/Ham's F-12 | 67% | Sigma | Cat#51651C |
| DMEM, high glucose | 33% | Sigma | Cat#D5796 |
| Fetal bovine serum | 5%(v/v) | Interpath | Cat#SFBSF15 |
| Penicillin-Streptomycin | 0.2%(v/v) | Life Tech | Cat#15070-063 |
| Hydrocortisone | 400ng/ml | Sigma | Cat#H0888 |
| Insulin | 5µg/ml | Sigma | Cat#I0516 |
| Cholera toxin | 8.4ng/ml | Sigma | Cat#C8052 |
| hEGF | 10ng/ml | Bioscientific | Cat#236-EG-200 |
| Adenine | 23.9µg/ml | Sigma | Cat#A2786 |
| Y-27632 | 10µM | Enzo life sciences | Cat#ALX-270-333-M025 |

**Supplementary Table S2.** Components of Lonza ALI base differentiation media.

| Components | Concentration | Supplier |
| --- | --- | --- |
| BEBM | 50% | Lonza, Cat#CC3171 |
| DMEM | 50% | Sigma, Cat#D5796 |
| Hydrocortisone | 0.1% | Lonza, Cat# (CC- 4031 F) |
| Insulin, Bovine | 0.1% | Lonza, Cat# (CC – 4021 F) |
| Epinephrine | 0.1% | Lonza, Cat# (CC – 4221 F) |
| Bovine Pituitary Extract | 0.4% | Lonza, Cat# (CC – 4009F) |
| Transferrin | 0.1% | Lonza, Cat# (CC – 4205F) |
| Ethanolamine | 80µM | Sigma, Cat#E0135 |
| hEGF | 10 ng/ml | Bioscientific Cat#236-EG-200 |
| All-trans retinoic acid | 30ng/ml | Sigma, Cat#R2625 |
| MgCl<sub>2</sub> | 0.3mM | Sigma, Cat#M8266 |
| MgSO<sub>4</sub> | 0.4mM | Sigma, Cat#M2643 |
| BSA | 0.5mg/ml | Sigma, Cat#A8806 |
| Penicillin/streptomycin | 2% | Life Tech, Cat#15070-063 |
| Amphotericin B solution | 250µg/ml | Sigma, Cat#A2942 |

**Supplementary Table S3.** Components of StemCell media expansion and differentiation.

| Components | Concentration | Supplier |
| --- | --- | --- |
| PneumaCult™ Ex-Plus media |  | StemCell, Cat#05041 |
| PneumaCult™ Ex-Plus media 50x supplement | Unknown | StemCell, Cat#05042 |
| PneumCult™ ALI maintenance media |  | StemCell, Cat#05002 |
| PneumaCult™-ALI 10x supplement | Unknown | StemCell, Cat#05003 |
| PneumaCult™-ALI maintenance supplement 100x | Unknown | StemCell, Cat#05006 |
| Hydrocortisone | 0.48µg/ml | StemCell, Cat#07925 |
| Heparin | 4µg/ml | StemCell, Cat#07980 |
| Penicillin/streptomycin | 2% | Life Tech, Cat#15070-063 |
| Amphotericin B solution | 250µg/ml | Sigma, Cat#A2942 |

**Supplementary Table S4.**
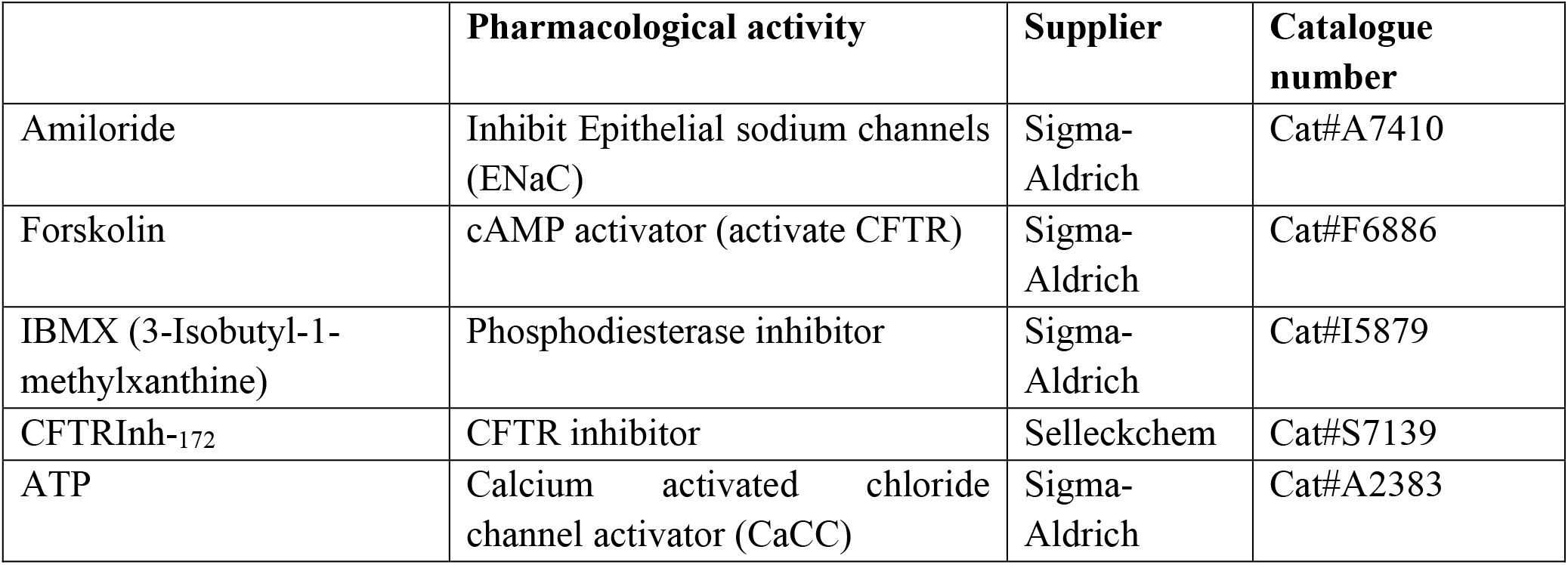
Pharmacologic compounds used for Ussing chamber measurements.

|  | Pharmacological activity | Supplier | Catalogue number |
| --- | --- | --- | --- |
| Amiloride | Inhibit Epithelial sodium channels (ENaC) | Sigma-Aldrich | Cat#A7410 |
| Forskolin | cAMP activator (activate CFTR) | Sigma-Aldrich | Cat#F6886 |
| IBMX (3-Isobutyl-1-methylxanthine) | Phosphodiesterase inhibitor | Sigma-Aldrich | Cat#I5879 |
| CFTRInh-172 | CFTR inhibitor | Selleckchem | Cat#S7139 |
| ATP | Calcium activated chloride channel activator (CaCC) | Sigma-Aldrich | Cat#A2383 |

**Supplementary Table S5.** Genes assessed by PCR.

| Gene | TaqMan Assay ID | Dye-Probe | Supplier | Catalogue number |
| --- | --- | --- | --- | --- |
| FOXJ1 | Hs00201755_m1 | FAM-MGB | ThermoFisher Scientific | 4331182 |
| SPDEF | Hs0017942_m1 | FAM-MGB | ThermoFisher Scientific | 4331182 |
| SCGB1 | Hs00171092_m1 | FAM-MGB | ThermoFisher Scientific | 4331182 |
| 18s | N/A | VIC-MGB | ThermoFisher Scientific | 4319413E |

**Supplementary Table S6.** Short-circuit currents and electrophysiologic parameters in ALI base differentiation media. Data represent short circuit current values for baseline currents and resistance of monolayers. Amiloride inhibited ENaC currents (ΔAmil), Forskolin/IBMX stimulated cAMP currents, CFTR_Inh-172_ inhibited currents (ΔFsk + CFTR_Inh-172_), and ATP-activated currents (ΔATP). Values represented are (mean ± SEM); D = Donor.

| AB-ALI | Baseline currents (μA/cm<sup>2</sup>) | Resistance ($\Omega$.cm<sup>2</sup>) | ΔAmil (μA/cm<sup>2</sup>) | ΔFsk/IBMX (μA/cm<sup>2</sup>) | ΔFsk/IBMX + CFTR<sub>Inh-172</sub> (μA/cm<sup>2</sup>) | ΔATP (μA/cm<sup>2</sup>) |
| --- | --- | --- | --- | --- | --- | --- |
| D1 | 17.76 ± 1.29 | 319.4 ± 25.99 | -6.29 ± 0.50 | 14.63 ± 2.28 | -22.11 ± 1.77 | 1.07 ± 0.35 |
| D2 | 22.30 ± 0.81 | 227.7 ± 14.24 | -3.20 ± 0.64 | 13.63 ± 2.02 | -23.97 ± 1.89 | 2.54 ± 0.27 |
| D3 | 23.20 ± 1.129 | 217.3 ± 25.47 | -12.21 ± 1.27 | 14.68 ± 0.89 | -24.07 ± 0.80 | 4.32 ± 0.30 |
| D4 | 17.50 ± 3.012 | 164.5 ± 29.21 | -7.93 ± 0.19 | 13.01±1.67 | -14.03 ± 1.48 | 2.38 ± 0.30 |
| D5 | 21.23 ± 2.03 | 140.5 ± 20.99 | -10.90 ± 0.34 | 5.17 ± 1.22 | -10.80 ± 0.50 | 1.12 ± 0.15 |

**Supplementary Table S7.** Short-circuit currents and electrophysiologic parameters in PneumaCult differentiation media. Data represent short circuit current values for baseline currents and resistance of monolayers. Amiloride inhibited ENaC currents (ΔAmil), Forskolin/IBMX stimulated cAMP currents, CFTR_Inh-172_ inhibited currents (ΔFsk + CFTR_Inh-172_), and ATP-activated currents (ΔATP). Values represented are (mean ± SEM); D = Donor.

| **PN**<br>**ALI** | **Baseline**<br>**currents**<br>**(μA/cm<sup>2</sup>)** | **Resistance**<br>**($\Omega$.cm<sup>2</sup>)** | **ΔAmil**<br>**(μA/cm<sup>2</sup>)** | **ΔFsk/IBMX**<br>**(μA/cm<sup>2</sup>)** | **ΔFsk/IBMX +**<br>**CFTR<sub>inh-172</sub>**<br>**(μA/cm<sup>2</sup>)** | **ΔATP**<br>**(μA/cm<sup>2</sup>)** |
| --- | --- | --- | --- | --- | --- | --- |
| D1 | - | - | - | - | - | - |
| D2 | 60.67 ± 11.23 | 88.77 ± 8.24 | -15.73 ± 7.46 | 12.07 ± 3.95 | -53.05 ± 6.48 | 7.74 ± 0.89 |
| D3 | 26.88 ± 3.84 | 69.10 ± 5.29 | -6.05 ± 1.26 | 7.47 ± 0.69 | -26.52 ± 4.43 | 7.90 ± 2.51 |
| D4 | 30.63 ± 1.018 | 117.6 ± 10.63 | -11.48 ± 0.68 | 22.40 ± 1.93 | -22.93 ± 2.84 | 7.28 ± 0.70 |
| D5 | 16.10 ± 0.71 | 211.0 ± 28.29 | -11.15 ± 0.74 | 7.78 ± 1.19 | -12.99 ± 1.20 | 3.44 ± 0.55 |

**Figure S1:**
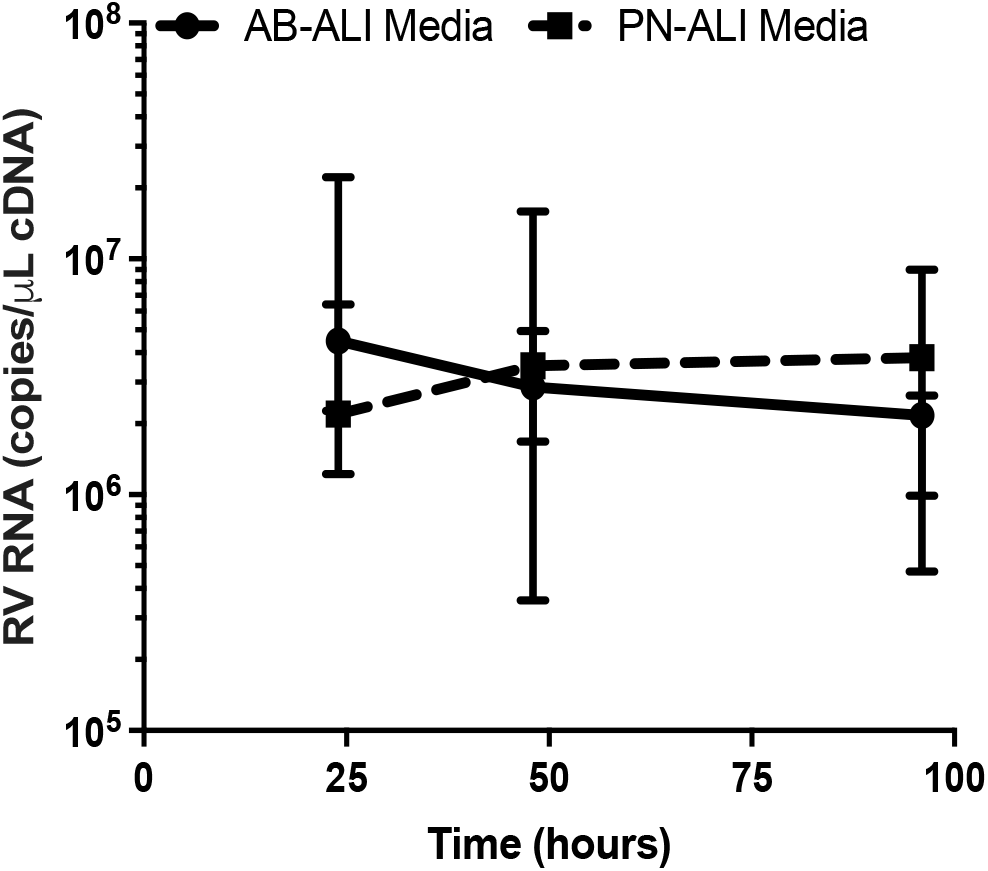
Line graph for viral growth kinetics in AB-ALI and PN-ALI CRpBECs.

## References

1. Kesimer M, Kirkham S, Pickles RJ, Henderson AG, Alexis NE, Demaria G, et al. Tracheobronchial air-liquid interface cell culture: a model for innate mucosal defense of the upper airways? Am J Physiol Lung Cell Mol Physiol. 2009 Jan;296(1):L92–100.

2. Pezzulo AA, Starner TD, Scheetz TE, Traver GL, Tilley AE, Harvey BG, et al. The air-liquid interface and use of primary cell cultures are important to recapitulate the transcriptional profile of in vivo airway epithelia. Am J Physiol Lung Cell Mol Physiol. 2011 Jan;300(1):L25–31.

3. Fulcher ML, Randell SH. Human Nasal and Tracheo-Bronchial Respiratory Epithelial Cell Culture. In: Randell SH, Fulcher ML, editors. Epithelial Cell Culture Protocols [Internet]. Totowa, NJ: Humana Press; 2012 [cited 2023 Mar 16]. p. 109–21. (Methods in Molecular Biology; vol. 945). Available from: https://link.springer.com/10.1007/978-1-62703-125-7_8

4. Suprynowicz FA, Upadhyay G, Krawczyk E, Kramer SC, Hebert JD, Liu X, et al. Conditionally reprogrammed cells represent a stem-like state of adult epithelial cells. Proc Natl Acad Sci U S A. 2012 Dec 4;109(49):20035–40.

5. Liu X, Ory V, Chapman S, Yuan H, Albanese C, Kallakury B, et al. ROCK inhibitor and feeder cells induce the conditional reprogramming of epithelial cells. Am J Pathol. 2012 Feb;180(2):599–607.

6. Reynolds SD, Rios C, Wesolowska-Andersen A, Zhuang Y, Pinter M, Happoldt C, et al. Airway Progenitor Clone Formation Is Enhanced by Y-27632-Dependent Changes in the Transcriptome. Am J Respir Cell Mol Biol. 2016 Sep;55(3):323–36.

7. Ruiz García S, Deprez M, Lebrigand K, Cavard A, Paquet A, Arguel MJ, et al. Novel dynamics of human mucociliary differentiation revealed by single-cell RNA sequencing of nasal epithelial cultures. Development. 2019 Oct 15;146(20):dev177428.

8. Broadbent L, Manzoor S, Zarcone MC, Barabas J, Shields MD, Saglani S, et al. Comparative primary paediatric nasal epithelial cell culture differentiation and RSV-induced cytopathogenesis following culture in two commercial media. Tripp RA, editor. PLOS ONE. 2020 Mar 26;15(3):e0228229.

9. Leung C, Wadsworth SJ, Yang SJ, Dorscheid DR. Structural and functional variations in human bronchial epithelial cells cultured in air-liquid interface using different growth media. Am J Physiol-Lung Cell Mol Physiol. 2020 May 1;318(5):L1063–73.

10. Luengen AE, Kniebs C, Buhl EM, Cornelissen CG, Schmitz-Rode T, Jockenhoevel S, et al. Choosing the Right Differentiation Medium to Develop Mucociliary Phenotype of Primary Nasal Epithelial Cells In Vitro. Sci Rep. 2020 Dec;10(1):6963.

11. Saint-Criq V, Delpiano L, Casement J, Onuora JC, Lin J, Gray MA. Choice of Differentiation Media Significantly Impacts Cell Lineage and Response to CFTR Modulators in Fully Differentiated Primary Cultures of Cystic Fibrosis Human Airway Epithelial Cells. Cells. 2020 Sep 21;9(9):E2137.

12. Livnat G, Meeker JD, Ostmann AJ, Strecker LM, Clancy JP, Brewington JJ. Phenotypic Alteration of an Established Human Airway Cell Line by Media Selection. Int J Mol Sci. 2023 Jan 8;24(2):1246.

13. Wark PAB, Johnston SL, Bucchieri F, Powell R, Puddicombe S, Laza-Stanca V, et al. Asthmatic bronchial epithelial cells have a deficient innate immune response to infection with rhinovirus. J Exp Med. 2005 Mar 21;201(6):937–47.

14. Awatade NT, Wong SL, Capraro A, Pandzic E, Slapetova I, Zhong L, et al. Significant functional differences in differentiated Conditionally Reprogrammed (CRC)- and Feeder-free Dual SMAD inhibited-expanded human nasal epithelial cells. J Cyst Fibros Off J Eur Cyst Fibros Soc. 2021 Mar;20(2):364–71.

15. Martinovich KM, Iosifidis T, Buckley AG, Looi K, Ling KM, Sutanto EN, et al. Conditionally reprogrammed primary airway epithelial cells maintain morphology, lineage and disease specific functional characteristics. Sci Rep. 2017 Dec 21;7(1):17971.

16. Reid AT, Nichol KS, Chander Veerati P, Moheimani F, Kicic A, Stick SM, et al. Blocking Notch3 Signaling Abolishes MUC5AC Production in Airway Epithelial Cells from Individuals with Asthma. Am J Respir Cell Mol Biol. 2020 Apr;62(4):513–23.

17. Smith CM, Djakow J, Free RC, Djakow P, Lonnen R, Williams G, et al. ciliaFA: a research tool for automated, high-throughput measurement of ciliary beat frequency using freely available software. Cilia. 2012 Dec;1(1):14.

18. Hsu ACY, See HV, Hansbro PM, Wark PAB. Innate immunity to influenza in chronic airways diseases. Respirol Carlton Vic. 2012 Nov;17(8):1166–75.

19. Hsu ACY, Dua K, Starkey MR, Haw TJ, Nair PM, Nichol K, et al. MicroRNA-125a and -b inhibit A20 and MAVS to promote inflammation and impair antiviral response in COPD. JCI Insight. 2017 Apr 6;2(7):e90443.

20. Johansen MD, Mahbub RM, Idrees S, Nguyen DH, Miemczyk S, Pathinayake P, et al. Increased SARS-CoV-2 Infection, Protease and Inflammatory Responses in COPD Primary Bronchial Epithelial Cells Defined with Single Cell RNA-Sequencing. Am J Respir Crit Care Med. 2022 May 12;

21. Lee DDH, Petris A, Hynds RE, O’Callaghan C. Ciliated Epithelial Cell Differentiation at Air-Liquid Interface Using Commercially Available Culture Media. In: Turksen K, editor. Epidermal Cells [Internet]. New York, NY: Springer US; 2019 [cited 2022 May 6]. p. 275–91. (Methods in Molecular Biology; vol. 2109). Available from: http://link.springer.com/10.1007/7651_2019_269

22. Sachs LA, Finkbeiner WE, Widdicombe JH. Effects of media on differentiation of cultured human tracheal epithelium. In Vitro Cell Dev Biol Anim. 2003 Feb;39(1–2):56–62.

23. Veerati PC, Nichol KS, Read JM, Bartlett NW, Wark PAB, Knight DA, et al. Conditionally reprogrammed asthmatic bronchial epithelial cells express lower FOXJ1 at terminal differentiation and lower IFNs following RV-A1 infection. Am J Physiol-Lung Cell Mol Physiol. 2022 Oct 1;323(4):L495–502.

24. Tosoni K, Cassidy D, Kerr B, Land SC, Mehta A. Using Drugs to Probe the Variability of Trans-Epithelial Airway Resistance. Vij N, editor. PLOS ONE. 2016 Feb 29;11(2):e0149550.

